# Adaptive regulation of dNTP homeostasis confers osimertinib resistance in EGFR mutant non-small cell lung carcinoma

**DOI:** 10.64898/2025.12.24.696437

**Authors:** Qian Xie, Yingying Wang, Yi Lei, Anthony Fernandez, Jessica D. Hess, Lin Yang, Terence Williams, Li Zheng, Binghui Shen, Min Li

**Author notes:** To whom correspondence should be addressed: Binghui Shen. These authors contribute to the work equally (Qian Xie, Yingying Wang).

## Abstract

Maintaining sustained deoxyribonucleotide triphosphate (dNTP) pools is essential for DNA replication fidelity and genome stability. In EGFR-mutant non-small cell lung carcinoma (NSCLC), we find that disruption of dNTP homeostasis plays a critical role in determining sensitivity to the EGFR inhibitor osimertinib and in shaping mechanisms of acquired resistance. Transcriptomic and biochemical analyses revealed that osimertinib suppresses RRM2 expression, a key regulator of dNTP synthesis, through the downregulation of the transcription factor MYBL2. In response to osimertinib-mediated replication stress and dNTP depletion, cells activated a compensatory pathway involving the stress-inducible ribonucleotide reductase subunit RRM2B via a novel transcriptional regulator, TNNT3. We further identified CHK2 signaling as essential for TNNT3 nuclear translocation and RRM2B transcriptional activation. Inhibition of CHK2 or combined CHK1/2 blockade impaired RRM2B induction, exacerbated replication stress, and delayed the development of osimertinib resistance both *in vitro* and in xenograft models. Collectively, our findings reveal that EGFR-mutant NSCLC cells rely on a dynamic EGFR–MYBL2–RRM2 and CHK2–TNNT3–RRM2B regulatory axis to maintain dNTP pool balance under therapeutic pressure. Disruption of this axis sensitizes tumors to osimertinib and impairs the acquisition of resistance, highlighting dNTP metabolism as a critical vulnerability and actionable target in EGFR-driven lung cancer.

**STATEMENT OF SIGNIFICANCE:** Targeting dNTP metabolism via the EGFR-MYBL2-RRM2 and CHK2-TNNT3-RRM2B regulatory axis sensitizes EGFR-mutant NSCLC to osimertinib treatment and delays drug resistance, revealing a novel and actionable therapeutic vulnerability in EGFR-driven lung cancer.

## INTRODUCTION

Uncontrolled proliferation is a hallmark of cancer, placing an extraordinary demand on DNA replication and repair machinery [1]. To meet this demand, cancer cells are highly dependent on nucleotide metabolism to maintain adequate pools of deoxyribonucleotide triphosphates (dNTPs), which are essential substrates for DNA synthesis and genome stability [2, 3]. Disruption of the dNTP pool balance, either through metabolic stress or therapeutic intervention, can impair replication fidelity, promote DNA damage, and drive genomic instability, ultimately contributing to tumor evolution and therapy resistance [4, 5].

Central to dNTP biosynthesis is ribonucleotide reductase (RNR), a highly regulated enzyme composed of the large catalytic subunit RRM1 and one of two small regulatory subunits, RRM2 or RRM2B. RNR is responsible for the reduction of 2’OH in ribonucleotides, converting them into deoxyribonucleotides. RRM2 is cell cycle-regulated and provides the bulk of dNTPs during S-phase, while RRM2B is induced under genotoxic or replication stress conditions and is thought to support DNA repair and replication restart [6–9]. Dysregulation of RNR subunit expression has been implicated in various cancers: elevated RRM2 is associated with aggressive tumor behavior and poor prognosis [10–13], while RRM2B plays context-dependent roles, acting as either a tumor suppressor to support DNA damage repair or a stress-adaptive factor depending on the oncogenic environment [14–17]. Importantly, recent studies have linked RNR subunit regulation to resistance mechanisms across multiple cancer therapies, including cytotoxic agents, radiation, and targeted inhibitors [18–21]. By altering dNTP supply in response to replication stress, cancer cells can adaptively reprogram nucleotide metabolism to preserve replication fork integrity and promote survival. This metabolic plasticity enables tumor cells to tolerate DNA damage, evade apoptosis, and acquire mutations that drive clonal evolution and therapeutic escape[19, 22].

In the context of EGFR-mutant non-small cell lung cancer (NSCLC), third-generation tyrosine kinase inhibitors (TKIs), such as osimertinib, initially induce strong clinical responses but are inevitably followed by acquired resistance. While several resistance mechanisms have been described—ranging from secondary EGFR mutations to the activation of bypass pathways [23–25]— less is known about how EGFR inhibition perturbs dNTP metabolism and whether RNR subunit rewiring contributes to therapy resistance. Clinical studies have linked RNR subunit expression with NSCLC progression and therapeutic outcome. High RRM2 expression correlates with tumor aggressiveness and poor prognosis [26–30], while RRM2B expression has been associated with both protective effects and adaptive resistance, depending on the treatment context [31, 32]. However, how EGFR signaling dynamically regulates RNR subunits in response to osimertinib, and how this contributes to resistance evolution, remains incompletely understood. Our work addresses this gap by identifying a previously unrecognized adaptive program in which TKI-mediated EGFR–MYBL2–RRM2 suppression is compensated by CHK2–TNNT3–RRM2B activation, allowing for restoration of the dNTP pool and sustained replication under EGFR-targeted therapy.

Here, we show that osimertinib disrupts the pyrimidine dNTP balance by downregulating RRM2, leading to replication stress and DNA damage. In response, cancer cells engage a CHK2-dependent transcriptional pathway involving TNNT3 to induce RRM2B, restoring dNTP pools and facilitating cell survival. Importantly, we demonstrate that pharmacologic inhibition of CHK2, or dual inhibition of CHK1/2, prevents RRM2B induction, aggravates replication stress, and significantly delays the development of osimertinib resistance *in vitro* and in cell-derived xenograft models. These findings underscore the functional significance of the CHK1/2 checkpoint pathways in not only sensing DNA damage, but also in sustaining adaptive survival under EGFR inhibition. By targeting CHK1/2 signaling, we reveal a strategy to disrupt dNTP recovery and replication resilience, sensitizing EGFR-mutant tumors to osimertinib and suppressing resistance acquisition. Together, our work identifies a previously unrecognized EGFR–MYBL2–RRM2 and CHK2–TNNT3–RRM2B regulatory network that preserves nucleotide homeostasis in osimertinib-treated NSCLC. These findings highlight dNTP metabolism and RNR subunit plasticity as central mediators of cancer cell adaptability, linking metabolic regulation to therapeutic resistance and disease progression.

## MATERIALS AND METHODS

### Cell culture and reagents

The PC-9 and HCC827 cell lines were obtained from ATCC and cultured in RPMI-1640 medium supplemented with 10% fetal bovine serum (FBS), penicillin (100 U/ mL), and streptomycin (100 μg/mL), at 37 °C and 5% CO2. The chemical compounds were purchased from MedChemExpress: MG132 (#HY-13259), Osimertinib (#HY-15772), PV-1019 (#HY-125203) and LY2606368 (#HY-18174). RRM2B-FLAG (#HG3150-CF) and MYBL2-HA (#HG14536-CY) plasmids were purchased from Sino Biological.

### Antibodies

The following antibodies and dilutions were used for Western Blot analysis. RRM1 (#10526-1-AP), RRM2 (11661-1-AP), RRM2B (#18005-1-AP), CHK1 (#25887-1-AP), CHK2 (#13954-1-AP), EGFR (#66455-1-Ig), beta Actin (#20536-1-AP), HA tag (#51064-2-AP), Histone H3 (#17168-1-AP), GAPDH (#60004-1-Ig), POLD1(#15646-1-AP), POLH (#28133-1-AP), MYBL2 (#18896-1-AP) and TNNT3 (#19729-1-AP) were purchased from Proteintech. Pol Ⅱ (8WG16) (#sc-56767) and Normal mouse IgG (#sc-2025) were purchased from Santa Cruz. Phospho-Chk1(ser345) (#133D3) and Phospho-Chk2 (Thr68) (#C13C1) were purchased from CST.

### Comet assay

Comet assays were carried out to analyze DNA damage using the CometAssay® Kit (R&D Systems) and according to a published protocol. Briefly, HCC827 and PC9 cells were seeded in 6-well plates at a density of 2×10^5^ cells/well with RPMI 1640 complete medium overnight and then treated with DMSO, osimertinib (10 nM), dNTP (25 µM)/siRRM2B, or a combination of dNTP/siRRM2B and osimertinib. After treatment for 6 or 24 hours, cells were collected in PBS and combined at a density of 1×10^5^ cells/mL with molten LMAgarose (37°C) at a ratio of 1:10 (v/v), followed by the immediate pipetting of 100 µL onto CometSlides^TM^. The slides were placed at 4 °C in the dark for 30 minutes and then immersed in lysis buffer at 4 °C for 1 hour. After removing excess buffer, slides were equilibrated in 1× TBE buffer at 4 °C for 15 minutes, followed by electrophoresis at 21 V for 40 minutes. Slides were then rinsed in deionized water for 5 minutes, fixed in 70% ethanol for 5 minutes, and air-dried at 37 °C for 15 minutes. For staining, 100 µL of diluted SYBR™ Gold (Invitrogen) was applied to each agarose spot and incubated for 30 minutes in the dark. Comet images were captured using a ZEISS Axio Observer fluorescence microscope, and DNA damage was quantified using Image J analysis software.

### DNA fiber assay

Cells were treated with osimertinib (1 μM), dNTP (25 μM), or a combination of these treatments. Cells were then trypsinized, embedded in agarose plugs (TopVision, Thermo Fisher Scientific), then lysed at 50 °C in a solution containing 1% n-lauroylsarcosine, 0.5 M EDTA (pH 8.0), and 0.2 mg/mL proteinase K for 72 hours. Plugs were then washed with TE buffer and proteinase K neutralized with 200 μM PMSF. The plugs were then melted and agarose was digested using β-agarase (NEB). Subsequently, the DNA was stretched on glass slides coated with 3-aminopropyltriethoxysilane (Sigma-Aldrich). DNA was denatured using 0.1 M NaOH in 70% Ethanol, then fixed with 0.5% glutaraldehyde. Slides were then blocked with 1% BSA for 20 min. DNA was stained using 1:100 mouse anti-IdU antibody (BD) and 1:500 rat anti-CldU (Abcam), followed by 1:100 Alexa Fluor 568 goat anti-mouse (Invitrogen) and 1:50 Dylight 488 goat anti-rat (Invitrogen). The stained DNA fibers were then imaged on a Zeiss Axio Observer Microscope.

### Drug escalation assay

To model the development of acquired drug resistance, a stepwise drug escalation assay was performed. HCC827 or PC9 cells were initially seeded at 30–40% confluence in standard growth medium (RPMI 1640 supplemented with 10% fetal bovine serum and 1% penicillin-streptomycin) and allowed to attach overnight. The following day, cells were exposed to 10 nM osimertinib, with parallel vehicle-treated cells serving as controls. Cells were maintained at 37 °C with 5% CO2 under continuous drug pressure and passaged upon reaching ∼80% confluence. At each passage, the drug concentration was doubled, provided that the majority of cells remained viable and proliferative. Starting at a drug concentration of 10 nM, this escalation process was continued over a period of 6–12 weeks until the cells exhibited stable growth at 1000 nM.

### RT-qPCR, RNA-seq, and analysis

Total RNA was extracted using TRIzol™ Reagent (Invitrogen) according to the manufacturer’s protocol. Briefly, cells were lysed in TRIzol, and phase separation was induced by the addition of chloroform. After centrifugation, the aqueous phase containing RNA was collected, and RNA was precipitated with isopropanol, washed with 75% ethanol, and resuspended in RNase-free water. RNA concentration and purity were determined using a NanoDrop spectrophotometer (Thermo Fisher Scientific). For cDNA synthesis, 1 µg of total RNA was reverse transcribed using the High-Capacity cDNA Reverse Transcription Kit (Applied Biosystems) in a 20 µL reaction volume with random primers, following the manufacturer’s instructions. Quantitative PCR (qPCR) was carried out using SYBR Green Master Mix (e.g., PowerUp SYBR Green, Applied Biosystems) on a QuantStudio Real-Time PCR System (Applied Biosystems). Each 20 µL reaction contained 1 µL of diluted cDNA, 10 µL of SYBR Green Master Mix, and 0.2 µM of forward and reverse primers. The cycling conditions were as follows: initial denaturation at 95 °C for 10 minutes, followed by 40 cycles of 95 °C for 15 seconds and 60 °C for 1 minute. A melt curve analysis was performed to confirm the specificity of amplification. Relative gene expression was calculated using the ΔΔCt method, normalized to GAPDH or ACTB as endogenous controls. All reactions were performed in technical triplicates, and each experiment was independently repeated at least three times. Primer sequences are available upon request or provided in **Supplementary Table S1**.

Sequencing libraries were generated using NEBNext® Ultra^TM^ RNA Library Prep Kit for Illumina (NEB, USA), following the manufacturer’s recommendations. The libraries were then subjected to next-generation sequencing on the Illumina HiSeq 2500 platform. The raw reads were quality-checked by FastQC v0.20.0 (https://www.bioinformatics.babraham.ac.uk/projects/fastqc/). The reads containing adapters, ploy-N, or any bases with Phred quality scores lower than 20 were removed with Cutadapt v2.1 and TrimGalore v0.6.5 (https://www.bioinformatics.babraham.ac.uk/projects/trim_galore/). The clean reads were aligned to the human genome (GRCh8) by HISAT2 v2.2.1. Read counts were calculated by featureCounts v2.0.6. and differential analysis of genes was performed using the R package DESeq2 (http://www.bioconductor.org/packages/release/bioc/html/DESeq2.html).

### Tumor growth and reduction in xenograft mice

All animal experiments were performed in accordance with a protocol approved by the Institutional Animal Care and Use Committee (IACUC) at City of Hope, following established institutional policies and procedures. Six- to eight-week-old male or female NSG (NOD SCID gamma) mice were subcutaneously implanted with 4 × 10⁶ HCC827 cells per mouse. Treatment began once tumor volumes reached approximately 400–500 mm³. Osimertinib was administered daily via oral gavage, and LY2606368 was administered via intraperitoneal injection. Tumor volumes were measured twice weekly using digital calipers and calculated using the formula: (Length × Width²) / 2 (mm³). Mice were euthanized when tumor volumes reached 1,000 mm³ or upon signs of morbidity, in accordance with humane endpoint criteria.

### Chromatin immunoprecipitation (ChIP)

ChIP assays were performed to evaluate the occupancy of RNA Polymerase II (RNAPII) and HA-tagged MYBL2 at target gene promoters, including RRM2. PC-9 and HCC827 cells were either left untransfected (for RNAPII ChIP) or transfected with a plasmid encoding HA-tagged MYBL2 using Lipofectamine 3000 (Thermo Fisher) according to the manufacturer’s protocol (for HA-ChIP). Forty-eight hours post-transfection, cells were crosslinked with 1% formaldehyde for 10 minutes at room temperature and quenched with 125 mM glycine for 5 minutes. Cells were washed with ice-cold PBS and lysed in SDS lysis buffer containing protease inhibitors. Chromatin was sheared by sonication using a Bioruptor (Diagenode) to generate DNA fragments of 200–500 bp. The lysates were cleared by centrifugation and pre-cleared with Protein A/G magnetic beads for 1 hour at 4 °C. For immunoprecipitation, samples were incubated overnight at 4 °C with either 5 µg of anti-RNAPII (8WG16) or anti-HA antibody. Normal mouse IgG served as a negative control. Immune complexes were captured with Protein A/G magnetic beads and sequentially washed with low-salt, high-salt, LiCl, and TE buffers. DNA-protein complexes were eluted and reverse crosslinked at 65 °C overnight, followed by RNase A and proteinase K treatment. DNA was purified using a spin column-based kit. Quantitative PCR (qPCR) was performed using SYBR Green Master Mix with primers targeting promoter regions of interest. ChIP enrichment was calculated as a percentage of input DNA and/or normalized to IgG controls. Each experiment was performed in triplicate and independently repeated at least three times.

### Colony formation assay

To assess long-term cell survival and proliferative capacity, colony formation assays were performed. PC-9 and HCC827 cells were transfected with the indicated siRNAs or plasmids and seeded into 6-well plates at a density of 500–1000 cells per well, depending on cell line growth rate. After 24 hours, cells were treated with varying concentrations of osimertinib with additional agents as specified. Cells were maintained under standard culture conditions (37 °C, 5% CO₂) for 10–14 days, with medium (containing drugs where applicable) refreshed every 3–4 days. Colonies were fixed with 4% paraformaldehyde for 15 minutes and stained with 0.5% crystal violet in 25% methanol for 30 minutes at room temperature. The plates were washed gently with water and then air-dried. Colonies consisting of ≥ 50 cells were counted manually or quantified using ImageJ software. Colony numbers were normalized to the untreated or control group and expressed as relative colony density. Each experiment was performed in triplicate and repeated independently at least three times.

### Cell viability assay using an ATP-based luminescence assay

Cell viability was assessed using the CellTiter-Glo® Luminescent Cell Viability Assay (Promega), which quantifies intracellular ATP levels as an indicator of metabolically active, viable cells. PC-9 and HCC827 cells were seeded in 96-well plates at a density of 1,000–2,000 cells per well in 100 µL of complete growth medium. After 24 hours, cells were treated with various concentrations of osimertinib or other indicated compounds for 72 hours. At the endpoint, 100 µL of CellTiter-Glo reagent was added directly to each well. The plates were then shaken for 2 minutes to induce cell lysis, followed by a 10-minute incubation at room temperature to stabilize the luminescent signal. Luminescence was measured using a microplate reader. Background signals from cell-free wells were subtracted, and data were normalized to vehicle-treated control wells. Results were expressed as a percentage of viable cells relative to control. All treatments were performed in triplicate wells, and each experiment was repeated independently at least three times.

### dNTPs Quantitation

Intracellular deoxynucleotide triphosphate (dNTP) levels were quantified using a fluorescence-based enzymatic assay, as previously described by Pai et al. [33], with minor modifications. Cells were washed twice with ice-cold PBS and lysed in 65% methanol to inactivate enzymes and extract dNTPs. Lysates were incubated at –80 °C for 1 hour, followed by centrifugation at 20,000 × g for 15 minutes at 4 °C. The supernatants were then collected and evaporated to dryness using a vacuum concentrator (SpeedVac). Dried extracts were resuspended in nuclease-free water and stored at – 80 °C until analysis. Quantification reactions were performed in 20 µL volumes containing: 1× Q5 Reaction Buffer (New England Biolabs), 0.25 µM primer, 0.25 µM long single-stranded DNA template specific for the target dNTP, 1× EvaGreen dye (Biotium), 20 U/mL Q5 High-Fidelity DNA Polymerase (New England Biolabs) and 1–2 µL of sample extract or dNTP standard. Each dNTP (dATP, dTTP, dCTP, dGTP) was measured in a separate reaction using template-primer pairs designed to make the reaction rate dependent on the limiting dNTP. Reactions were set up in 96-well plates and run on a real-time PCR instrument using the following thermal cycling conditions: 98 °C for 30 seconds, followed by 40 cycles of 98 °C for 10 seconds and 65 °C for 30 seconds.

Fluorescence was measured in real time, and the signal generated was proportional to the amount of incorporated dNTP. Standard curves were generated from serial dilutions of known concentrations of commercial dNTPs (Thermo Fisher Scientific), and sample concentrations were interpolated from the standard curves. dNTP levels were normalized to total cell number or total protein content.

### Cell fractionation

Nuclear and cytoplasmic protein fractions were prepared using a modified detergent-based method. Cells were harvested by trypsinization, washed twice with ice-cold PBS, and resuspended in ice-cold hypotonic lysis buffer (10 mM HEPES pH 7.9, 10 mM KCl, 1.5 mM MgCl₂, 0.5 mM DTT, 0.1% NP-40, and protease/phosphatase inhibitors). The cell suspension was incubated on ice for 10 minutes to allow swelling and lysis of the plasma membrane. Samples were then centrifuged at 800 × g for 5 minutes at 4 °C to pellet the nuclei. The supernatant was collected and clarified by centrifugation at 16,000 × g for 10 minutes. The nuclear pellet was washed once with hypotonic buffer and then resuspended in nuclear extraction buffer (20 mM HEPES pH 7.9, 400 mM NaCl, 1 mM EDTA, 1 mM EGTA, 1 mM DTT, 10% glycerol, and protease/phosphatase inhibitors). Samples were vortexed briefly and incubated on ice for 30 minutes with intermittent mixing to solubilize nuclear proteins. After centrifugation at 16,000 × g for 10 minutes at 4 °C, the supernatant containing nuclear proteins was collected. Protein concentrations were determined using the BCA assay, and equal amounts were used for downstream western blot analysis. GAPDH and Histone H3 were used as markers for the cytoplasmic and nuclear fractions, respectively, to confirm fractionation quality.

### Statistical analysis

All experiments were performed in biological triplicates unless otherwise stated. Quantitative data are presented as mean ± standard error of the mean (SEM) or standard deviation (SD) as indicated. Statistical analyses were conducted using GraphPad Prism. Comparisons between two groups were made using an unpaired two-tailed Student’s *t*-test, while comparisons involving more than two groups were analyzed using one-way or two-way ANOVA followed by appropriate post hoc tests (e.g., Tukey’s or Bonferroni correction). For survival analyses, Kaplan–Meier curves were plotted and significance assessed using the log-rank (Mantel–Cox) test. p values < 0.05 were considered statistically significant and are denoted as follows: * p < 0.05, ** p < 0.01, *** p < 0.001. For qPCR and ChIP-qPCR, fold enrichment or expression levels were calculated using the ΔΔCt method, normalized to housekeeping genes (e.g., GAPDH or ACTB), and compared to control conditions. ChIP results were expressed as either percent input or fold enrichment over IgG as specified in figure legends.

### Data Availability

The data generated in this study are available upon request from the corresponding author.

## RESULTS

### Osimertinib Treatment Alters Expression of DNA Replication and Repair Genes in EGFR-Mutant NSCLC Cells

To characterize the transcriptional response of EGFR-mutant non-small cell lung cancer (NSCLC) cells to the third-generation EGFR tyrosine kinase inhibitor, osimertinib, we performed RNA sequencing (RNA-seq) on PC-9 cells treated with osimertinib versus DMSO control. Differential expression analysis revealed a distinct set of genes that were significantly downregulated upon osimertinib exposure, many of which encode factors critical for DNA synthesis and cell cycle progression. In the volcano plot (**Fig. 1A**), transcripts such as RRM2, MYBL2, KPNA2, CCDN1, and MCM6 were among the top downregulated genes, whereas a smaller subset of genes (e.g., DEPP1, RND1, VTCN1) were significantly upregulated. Gene Ontology (GO) and KEGG pathway enrichment analysis of all significantly dysregulated genes demonstrated that pathways governing cell cycle progression, DNA replication, and DNA repair were among the most highly overrepresented categories following osimertinib treatment (**Fig. 1B**). In particular, the “Cell cycle” pathway exhibited the highest gene ratio and lowest false discovery rate, followed by “DNA replication”, “Base excision repair”, and “Homologous recombination”. Smaller but significant enrichment was noted for “Nucleotide metabolism” and “Pyrimidine metabolism,” indicating broad suppression of DNA maintenance programs and nucleotide synthesis.

**Figure 1.**
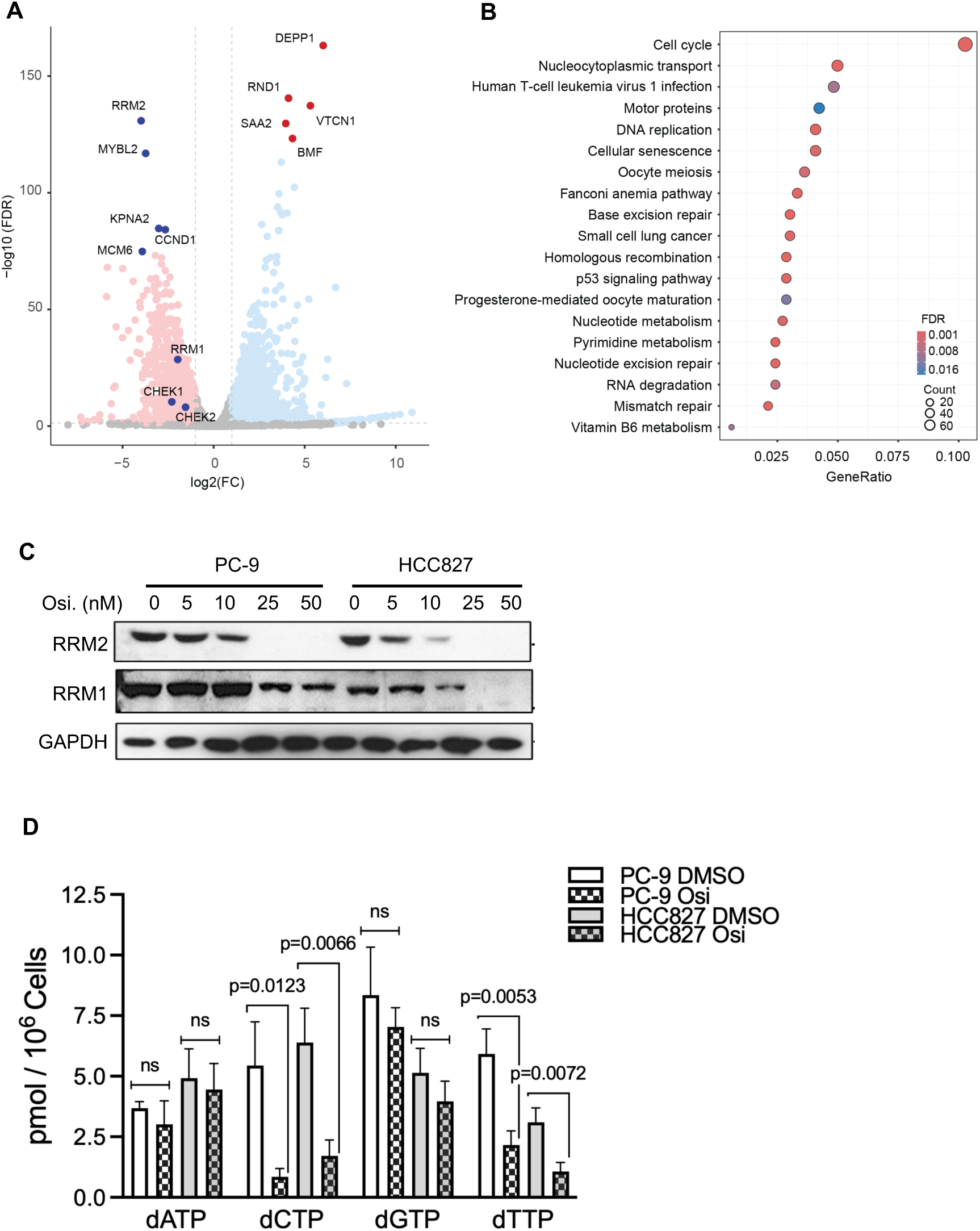
Osimertinib suppresses RRM2 expression and reduces dNTP levels in PC-9 and HCC827 cells. **(A)** Volcano plot showing gene expression changes in PC-9 cells after treatment with osimertinib (Osi). The x-axis denotes fold expression change, and the y-axis shows p-value of the change. Genes with a log₂ fold expression change ≥ 1 and an adjusted p-value < 0.05 are labeled. **(B)** KEGG pathway enrichment analysis of genes downregulated by Osi in PC-9 cells. Pathways are ranked based on –log₁₀ (p-value). **(C)** PC-9 and HCC827 cells were treated with the indicated concentrations of osimertinib for 24 hrs. Protein expression of RRM1 and RRM2 was assessed by western blot, with GAPDH as a loading control. Results shown are representative of three independent experiments. **(D)** Intracellular dNTP levels in PC-9 and HCC827 cells were quantified with or without 24-hour Osi treatment. Data are presented as mean ± S.D. from three independent biological replicates.

To validate that mRNA downregulation translated to reduced protein expression, we conducted western blot (WB) analysis of RRM2 in PC-9 and HCC827 cells treated with increasing concentrations of osimertinib for 24 hours. As shown in **Fig. 1C**, RRM2 protein levels decreased by over 70% in HCC827 cells and approximately 50% in PC-9 cells following 10 nM osimertinib treatment, relative to DMSO controls. In contrast, RRM1 levels remained unchanged at this dose, though excess treatment did result in lower levels. These results corroborate the RNA-seq data and confirm that osimertinib selectively suppresses RRM2 expression. Given the observed downregulation of RRM2, we next quantified the effect this had on intracellular deoxynucleotide triphosphate (dNTP) pools in PC-9 and HCC827 cells treated with osimertinib for 24 hours. As shown in **Fig. 1D**, osimertinib treatment did not significantly alter levels of dATP or dGTP in either cell line. In contrast, both dCTP and dTTP were markedly reduced in the Osi-treated groups: dCTP levels decreased from 6.3 ± 0.5 (PC-9 DMSO) to 1.2 ± 0.2 pmol/10⁶ cells, and from 5.8 ± 0.4 (HCC827 DMSO) to 1.0 ± 0.3 pmol/10⁶ cells. Similarly, dTTP levels dropped from 5.5 ± 0.6 to 1.3 ± 0.2 pmol/10⁶ cells in PC-9 cells (p < 0.01) and from 4.7 ± 0.3 to 0.9 ± 0.1 pmol/10⁶ cells in HCC827 cells (p < 0.01). Collectively, osimertinib treatment in EGFR-mutant NSCLC cell lines induces a transcriptional alteration that suppresses key DNA replication and repair genes (notably RRM2), disrupts cell cycle and DNA maintenance pathways, and leads to the selective depletion of pyrimidine dNTP pools (dCTP and dTTP), as shown in previous studies targeting RRM2 [34, 35], indicating that the salvage pathway for pyrimidine is less efficient compared to those for other deoxynucleotides. These data suggest that osimertinib’s cytotoxicity may, in part, derive from impairing nucleotide synthesis and destabilizing DNA replication fidelity.

### MYBL2 directly regulates RRM2 transcription in EGFR-mutant NSCLC cells

To investigate the transcriptional regulation of RRM2 in EGFR-mutant non-small cell lung cancer (NSCLC), we examined the role of the transcription factor MYBL2, which we previously found to be downregulated due to a lack of activation following osimertinib treatment. Prior studies have shown that MYBL2 can directly bind to the RRM2 promoter and activate its transcription during the S-phase in colorectal cancer cell lines [10]. Based on this, we first performed RNA polymerase II (RNAPII) chromatin immunoprecipitation (ChIP) assays to evaluate RNAPII recruitment to two predicted RRM2 promoter regions (PR1 and PR2) in the presence or absence of MYBL2. We found that MYBL2 knockdown significantly reduced RNAPII occupancy at both PR1 and PR2 in PC-9 and HCC827 cells, with the most pronounced reduction at PR2 (**Fig. 2A**). These findings indicate that MYBL2 promotes RNAPII recruitment to the RRM2 promoter. To evaluate MYBL2’s role in regulating the expression of RRM2, we performed western blot and RT-qPCR analyses and found a marked decrease in RRM2 protein and mRNA levels following MYBL2 knockdown in both cell lines, corresponding with reduced MYBL2 expression (**Supplementary Fig. S1A and B**). To determine whether osimertinib disrupts MYBL2-dependent transcription of RRM2, we conducted ChIP assays in PC-9 and HCC827 cells overexpressing HA-tagged MYBL2, with or without osimertinib treatment. MYBL2 overexpression significantly enhanced promoter occupancy at both PR1 and PR2 compared to control (**Fig. 2B**). However, osimertinib treatment markedly reduced MYBL2 binding PR2 in the context of overexpression. These results suggest that osimertinib impairs MYBL2’s ability to engage the RRM2 promoter, thereby inhibiting its transcriptional activation.

**Figure 2.**
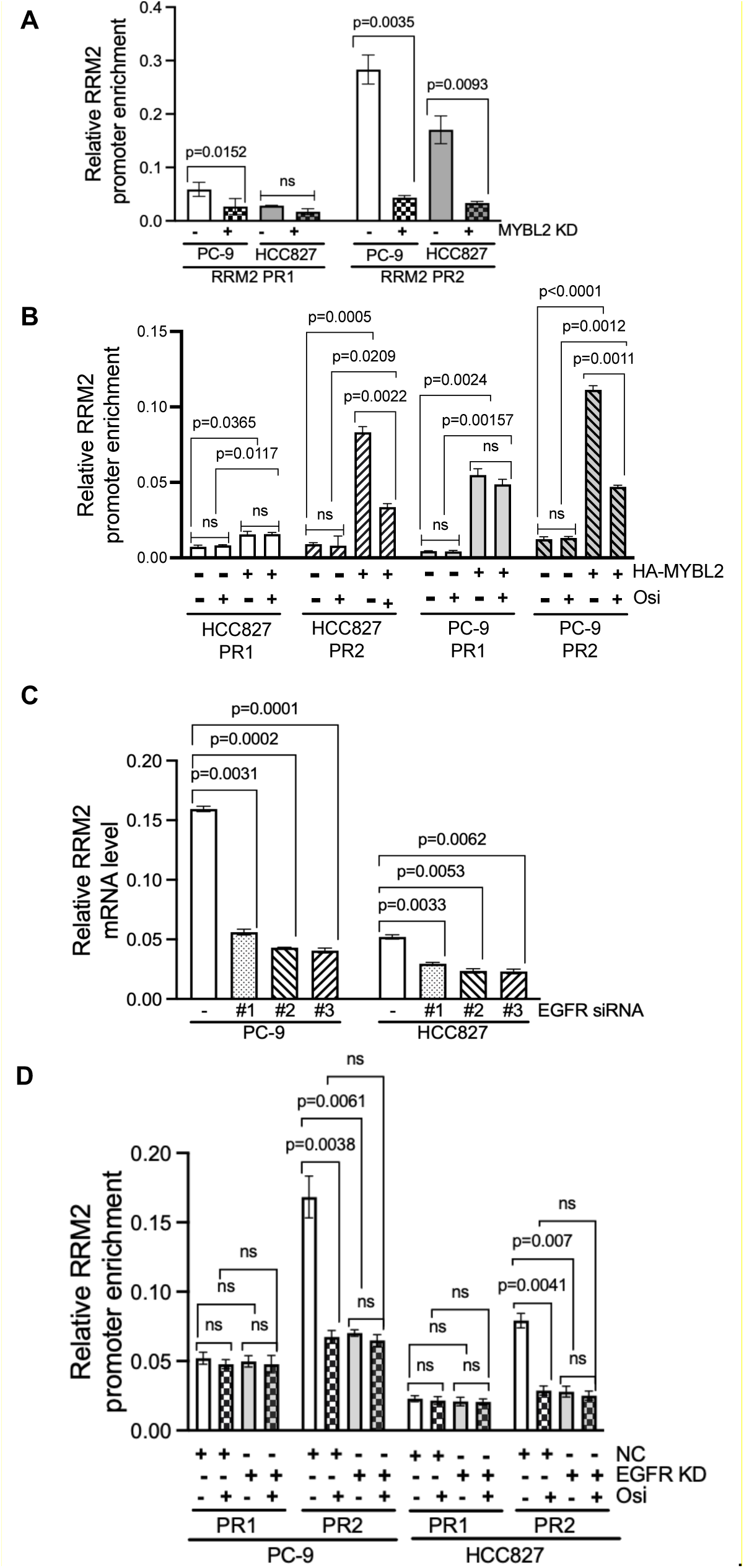
EGFR signaling promotes RRM2 transactivation through MYBL2 in PC-9 and HCC827 cells. **(A)** Chromatin immunoprecipitation followed by qPCR (ChIP-qPCR) was performed using an anti-RNAPII (8WG16) antibody to evaluate recruitment of RNA polymerase II to the RRM2 promoter, with or without osimertinib (Osi) treatment. Relative enrichment was normalized to the ACTB promoter. **(B)** ChIP-qPCR using HA-tagged MYBL2-overexpressing cell lines was conducted to assess MYBL2 binding to the RRM2 promoter. Promoter occupancy is shown relative to input. Data are presented as mean ± s.e.m. (n = 3). ***p < 0.001. **(C)** RRM2 mRNA expression was quantified by RT-qPCR in PC-9 and HCC827 cells following EGFR knockdown using siRNA. Transcript levels were normalized to GAPDH. **(D)** The impact of EGFR knockdown on MYBL2 recruitment to the RRM2 promoter was evaluated by ChIP-qPCR in HA-MYBL2-expressing cells transfected with control or EGFR siRNA. Promoter enrichment was normalized to input. Data are shown as mean ± s.e.m. (n = 3). ***p < 0.001. All data are representative of at least three independent experiments.

Given that MYBL2 transactivation is regulated by EGFR signaling, we next examined whether EGFR knockdown affects RRM2 mRNA expression. Using three independent siRNAs targeting EGFR in PC-9 and HCC827 cells, we observed a significant reduction in RRM2 mRNA levels following EGFR depletion, coinciding with efficient knockdown of EGFR protein (**Supplementary Fig. S1C; Fig. 2C**). These results suggest that EGFR signaling positively regulates RRM2 expression, likely through MYBL2-dependent transcriptional control. To determine whether EGFR modulates MYBL2’s recruitment to the RRM2 promoter, we performed ChIP-qPCR assays. EGFR knockdown significantly reduced MYBL2 enrichment at the RRM2 PR2 promoter site (**Fig. 2D**), and this effect was not further enhanced by osimertinib treatment in the EGFR-depleted background. However, in cells with intact EGFR expression, osimertinib treatment alone significantly reduced MYBL2 binding to the RRM2 promoter. These findings demonstrate that MYBL2 directly binds to and activates the RRM2 promoter, and that this activity is regulated by EGFR signaling.

Consistent with MYBL2’s essential role in regulating RRM2 transcription, its expression level also correlates with sensitivity to osimertinib. As shown in **Supplementary Fig. S2A**, HCC827 cells with MYBL2 knockdown were more sensitive to osimertinib compared to control cells, while MYBL2 overexpression conferred resistance. Similar results were observed in PC-9 cells (**Supplementary Fig. S2B**), where MYBL2 overexpression shifted the dose–response curve to the right, indicating reduced drug sensitivity, whereas MYBL2 knockdown increased sensitivity to osimertinib. These data support a model in which EGFR–MYBL2–RRM2 signaling is a key regulatory axis influencing osimertinib response in EGFR-mutant NSCLC.

### Osimertinib Impairs DNA Replication Fork Progression in EGFR-Mutant NSCLC Cells

To assess the effects of osimertinib on DNA replication dynamics, we performed DNA fiber assays in PC-9 and HCC827 cells treated with DMSO, Osi, exogenous dNTPs, or a combination of Osi and dNTPs. Cells were sequentially labeled with the thymidine analogs IdU (red) and CldU (green) with treatment to track nascent DNA synthesis before and after treatment. Representative DNA fiber images are shown in the upper panel (**Fig. 3A**), with red and green segments corresponding to IdU and CldU incorporation, respectively. In DMSO-treated cells, long, continuous red-green tracks were observed, indicative of normal replication fork progression. Osi-treated cells, however, displayed visibly shorter green tracks, reflecting impaired elongation. Interestingly, cells treated with dNTPs alone showed longer green tracks than DMSO controls, suggesting enhanced fork progression. The combination of osimertinib and dNTPs resulted in the longest green tracks, indicating a striking rescue and possible overcompensation of DNA synthesis.

**Figure 3.**
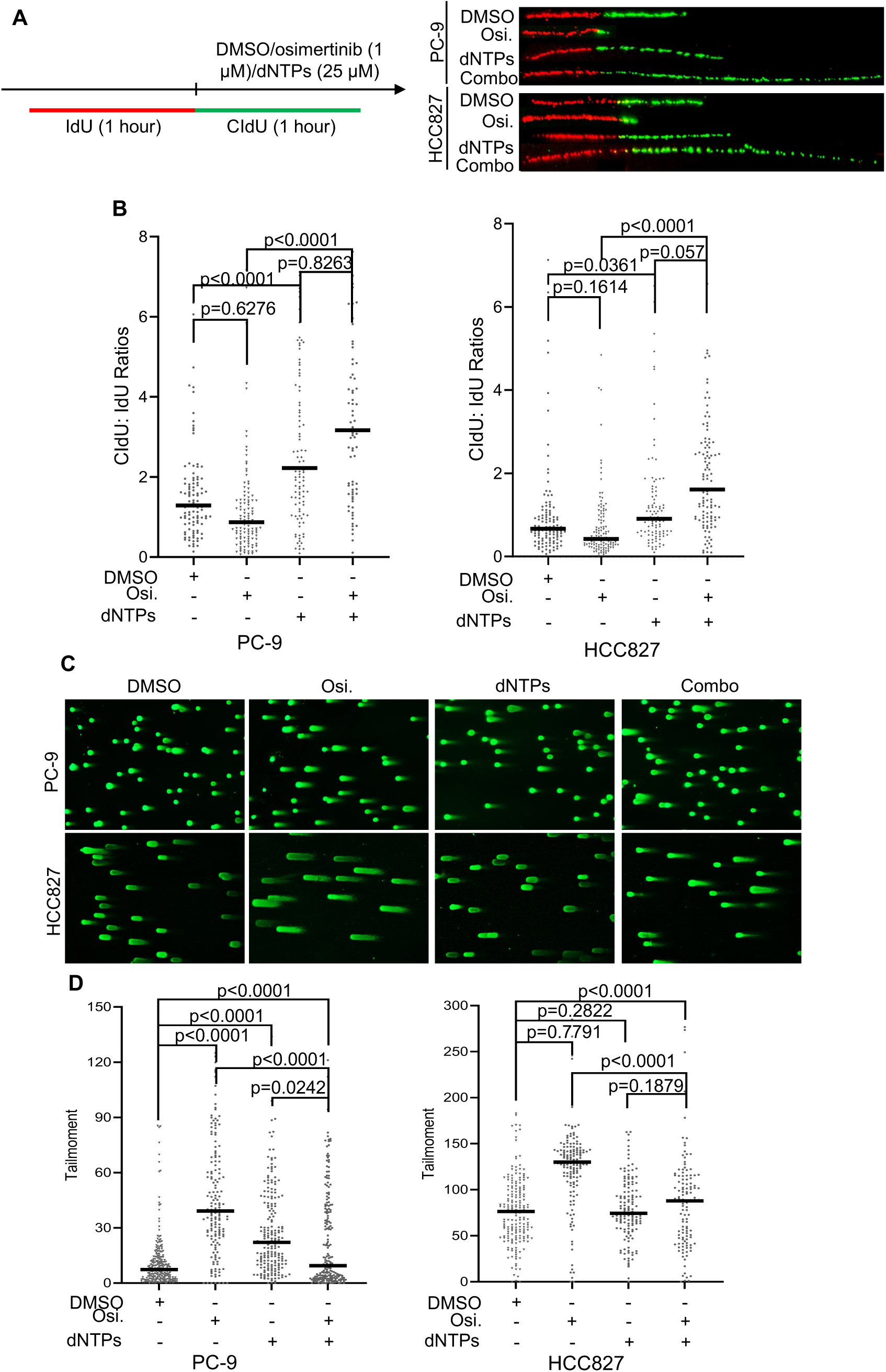
Osimertinib disrupts DNA replication fork progression and induces double-strand DNA breaks in EGFR-mutant NSCLC cells. **(A)** Representative DNA fiber images from the indicated cell lines under different treatment conditions. Cells were labeled sequentially with IdU (red) for 1 hour without treatment, followed by CldU (green) in the presence of osimertinib, dNTP supplementation, or their combination. DNA fibers were then isolated and spread to analyze replication dynamics. **(B)** Quantification of replication fork progression, shown as the ratio of CldU to IdU track lengths for individual DNA fibers. **(C)** Representative comet assay images showing DNA damage in HCC827 and PC-9 cells after 24-hour treatment with DMSO (control), osimertinib (5 µM), dNTPs (25 µM), or the combination of osimertinib and dNTPs. **(D)** Quantification of DNA damage, expressed as comet tail moment, under the indicated conditions. Data reflect the extent of double-strand DNA breaks in response to treatment.

Quantitative analysis of the CldU/IdU tract length ratio (**Fig. 3B**) revealed a significant reduction in CldU track length in Osi-treated cells compared to controls in both cell lines, confirming that osimertinib impairs fork progression. Exogenous dNTP supplementation alone accelerated replication, consistent with the known role of dNTP availability in regulating DNA polymerase activity. Unexpectedly, the combination of osimertinib and dNTPs caused an exceptional increase in CldU track length, suggesting aberrant or dysregulated DNA synthesis. Supporting this observation, WB analysis showed a modest increase in POLH levels in the chromatin fraction upon Osi/dNTPs co-treatment (**Supplementary Fig. S3**), implying that DNA polymerase switching in response to osimertinib-induced replication stress may contribute to the abnormal elongation phenotype observed. Together, these data demonstrate that osimertinib induces transcriptional downregulation of RRM2 and depletion of nucleotide pools, contributing to the disruption of DNA replication dynamics in EGFR-mutant cancer cell lines.

To further determine whether osimertinib-induced suppression of DNA replication leads to DNA damage in EGFR-mutant NSCLC cells, we performed neutral Comet assays in PC-9 and HCC827 cells. Cells were treated with DMSO, osimertinib, exogenous dNTPs, or a combination of osimertinib and dNTPs. Representative images (**Fig. 3C**) show that DMSO-treated cells displayed minimal comet tails, consistent with low baseline levels of double-strand DNA breaks (DSBs). In contrast, osimertinib-treated cells exhibited significantly greater tail moments, indicating increased DNA damage. Importantly, supplementation with dNTPs markedly reduced comet tail length and intensity in osimertinib-treated cells, suggesting that osimertinib-induced dNTP depletion contributes to replication stress and subsequent DNA breakage. Quantification of tail moments (**Fig. 3D**) confirmed these findings: Osimertinib significantly increased DNA damage in both PC-9 and HCC827 cells, whereas co-treatment with dNTPs partially rescued this effect, resulting in a notable reduction in tail moment. These results indicate that osimertinib-induced DNA damage is, at least in part, mediated by dNTP pool imbalance, and that replenishment of dNTPs can mitigate replication-associated DNA breaks.

### Induction of RRM2B Expression Following Osimertinib Treatment Is Essential for Restoring dNTP Pools and Minimizing DNA Breaks

Given the critical role of balanced dNTP pools in supporting DNA replication and repair, compensating for RRM2 loss through the activation of alternative pathways becomes essential for the survival of osimertinib-treated cells. To investigate the temporal regulation of ribonucleotide reductase subunits in response to osimertinib, we analyzed the protein and mRNA levels of RRM2 and its homolog RRM2B in PC-9 and HCC827 cells. WB analysis revealed a rapid and sustained reduction in RRM2 protein expression following osimertinib treatment, while RRM2B protein levels progressively increased over time in both cell lines (**Fig. 4A**). Consistent with these findings, RT-qPCR showed a significant decrease in RRM2 mRNA levels, accompanied by a steady upregulation of RRM2B transcripts, reaching a 3- to 5-fold increase by 48 hours post-treatment (**Fig. 4B**). These results indicate that RRM2B is transcriptionally induced as a compensatory mechanism to support dNTP synthesis in response to RRM2 suppression and replication stress.

**Figure 4.**
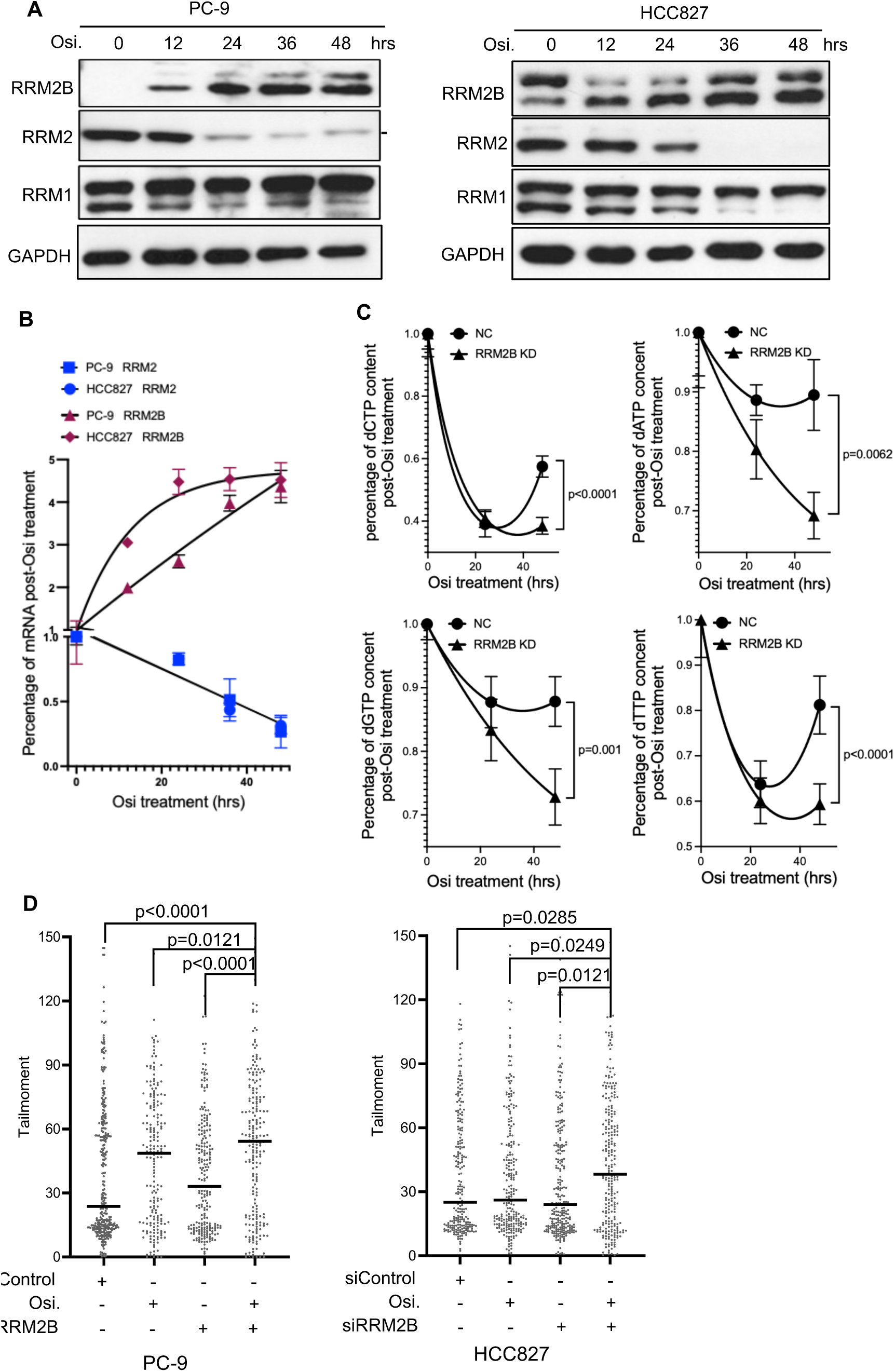
RRM2B induction is critical for replenishing dNTP pools and limiting DNA damage. **(A)** PC-9 and HCC827 cells were treated with 5 nM osimertinib (Osi) for the indicated durations. Protein levels of RRM1, RRM2, and RRM2B were assessed by western blotting, with GAPDH used as a loading control. Results are representative of three independent experiments. **(B)** Time-course analysis of RRM2 and RRM2B mRNA expression in PC-9 and HCC827 cells treated with 5 nM Osi. Transcript levels were measured by RT-qPCR, normalized to GAPDH, and expressed relative to untreated controls. Data represent three independent experiments. **(C)** Intracellular dNTP levels in PC-9 cells with or without RRM2B knockdown following 0, 24, or 48 hours of Osi treatment. dNTP levels were normalized to untreated controls. Data shown are representative of three independent experiments. **(D)** Quantification of DNA damage, expressed as comet tail moment, from comet assays in HCC827 and PC-9 cells treated with DMSO (control), osimertinib (10 nM), siRRM2B, or the combination of osimertinib (5 µM) and siRRM2B for 24 hours.

To investigate the functional significance of RRM2B induction during EGFR inhibition, we employed two independent siRNAs to knock down RRM2B in PC-9 cells. Knockdown efficiency was confirmed by a marked reduction in RRM2B mRNA levels following osimertinib treatment (**Supplementary Fig. S4A and B**). We next quantified intracellular dNTP pools under various treatment conditions to assess the impact on nucleotide metabolism. As shown in **Fig. 4C**, osimertinib treatment for 24 hours led to a substantial decrease in pyrimidine dNTPs in control siRNA-transfected cells, with dCTP and dTTP levels falling to approximately 40% and 65% of untreated levels, respectively. By 48 hours, a partial recovery was observed, with dCTP and dTTP levels rising to ∼60% and ∼80% of baseline, suggesting a delayed compensatory response.

In contrast, RRM2B-depleted cells exhibited a more profound and sustained suppression of pyrimidine dNTPs. At 48 hours, dCTP and dTTP levels remained significantly reduced at ∼40% and ∼60%, respectively, with no evidence of recovery. Notably, dATP and dGTP levels were also markedly reduced in RRM2B knockdown cells at this time point, consistent with previous reports indicating that RRM2B is essential for maintaining purine dNTP homeostasis [36]. These findings suggest that RRM2B plays a critical role in supporting both purine and pyrimidine dNTP synthesis under conditions of EGFR inhibition. Its induction during osimertinib treatment appears to be necessary for restoring dNTP pools, thereby providing a metabolic advantage that helps sustain DNA replication and promote cell survival in the face of EGFR pathway inhibition.

To assess whether RRM2B depletion exacerbates DNA damage under these conditions, we performed neutral Comet assays in PC-9 and HCC827 cells with or without RRM2B knockdown. As shown in **Supplementary Fig. S5** and **Fig. 4D**, osimertinib-treated control cells exhibited moderate DNA damage, as indicated by comet tail formation. However, RRM2B-depleted cells treated with osimertinib displayed markedly longer comet tails, indicative of increased double-strand breaks (DSBs). Quantification of tail moments confirmed a significant increase in DNA damage in RRM2B knockdown cells compared to control cells. These results highlight the critical role of RRM2B in maintaining nucleotide balance and genome integrity during EGFR inhibition. While osimertinib downregulates the canonical RRM2-driven nucleotide synthesis pathway, the induction of RRM2B acts as a compensatory mechanism that partially restores dNTP pools and mitigates replication-associated DNA damage. Disruption of this backup pathway sensitizes cells to EGFR-targeted therapy and contributes to therapeutic vulnerability.

### TNNT3 Drives RRM2B Transactivation to Promote Osimertinib Resistance

RRM2B was originally identified as a downstream target of p53, with its expression tightly regulated by p53 activity [7]. However, more recent studies have uncovered an unexpected regulatory role for troponin T3 (TNNT3) in controlling RRM2B expression in skeletal muscle tissue, independent of p53 [37]. Given that PC-9 and HCC827 cells lack wild-type p53, we hypothesized that these cells may employ TNNT3 as an alternative transcriptional regulator to drive RRM2B transactivation in response to osimertinib. To investigate this, we performed cellular fractionation followed by western blot analysis to determine whether TNNT3 translocates to the nucleus following osimertinib treatment. As shown in **Fig. 5A**, osimertinib treatment for 24 hours led to a notable increase in nuclear TNNT3 levels in both PC-9 and HCC827 cells, coinciding with the upregulation of RRM2B protein expression. To assess the functional relevance of TNNT3 in this context, we knocked down TNNT3 using two independent siRNAs. As shown in **Fig. 5B**, TNNT3 depletion markedly reduced RRM2B protein levels following osimertinib treatment. Consistent with these results, RT-qPCR analysis revealed that osimertinib-induced RRM2B mRNA expression was significantly blunted in TNNT3-depleted cells, whereas robust induction was observed in control siRNA-transfected cells (**Fig. 5C**). Together, these findings demonstrate that TNNT3 is essential for RRM2B transactivation in response to osimertinib and acts as a compensatory transcriptional regulator in the absence of functional p53 in EGFR-mutant NSCLC cells.

**Figure 5.**
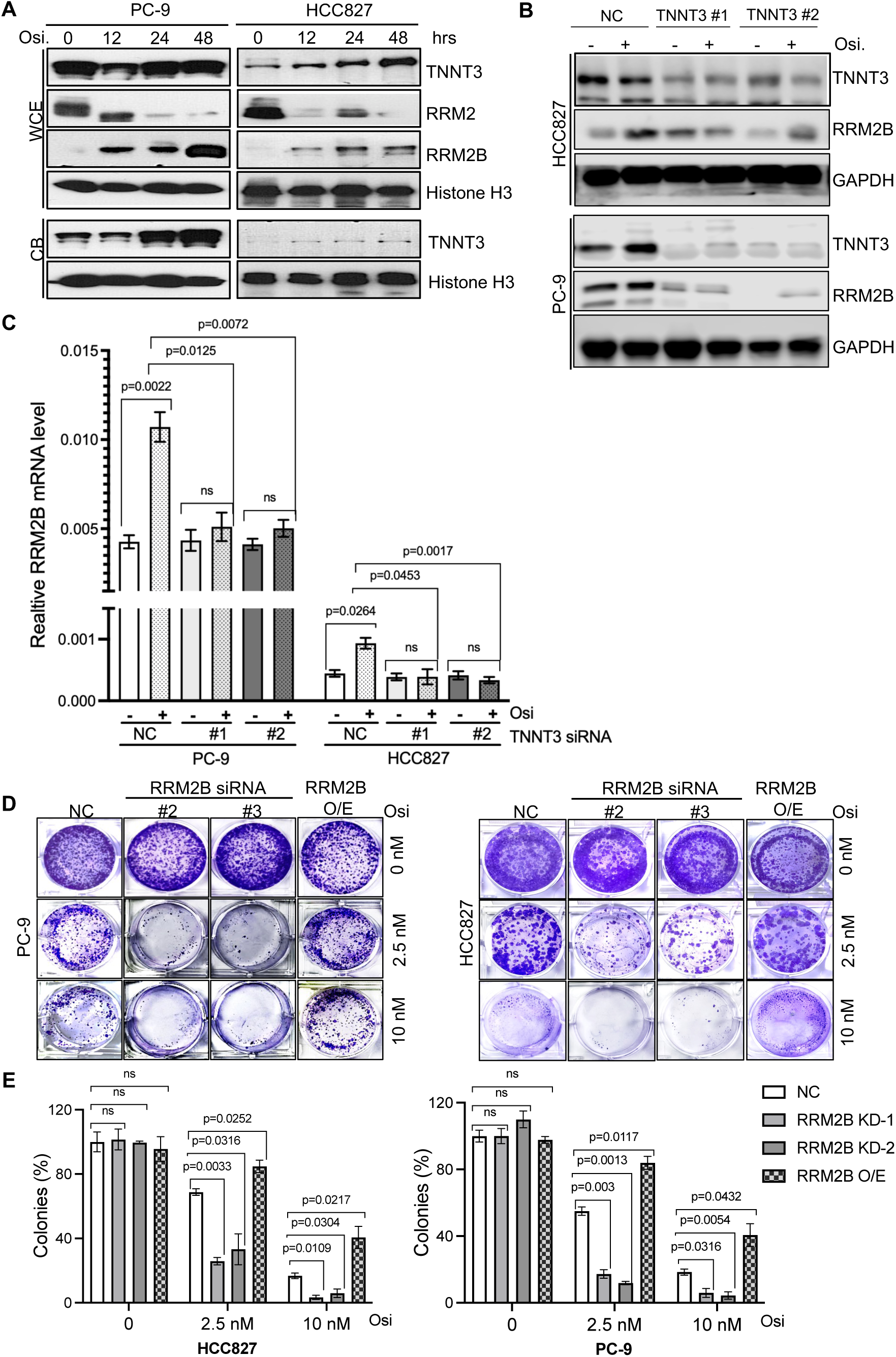
TNNT3-mediated transactivation of RRM2B contributes to osimertinib resistance. **(A)** PC-9 and HCC827 cells were treated with 5 nM osimertinib (Osi) for the indicated time points, and nuclear extracts were prepared. TNNT3 protein levels were analyzed by western blotting; ACTB and Histone H3 were used as loading controls. Data shown are representative of three independent experiments. **(B)** PC-9 and HCC827 cells were transfected with control or TNNT3 siRNA 24 hours prior to Osi treatment. Protein levels of TNNT3 and RRM2B were analyzed by western blotting, with ACTB as a loading control. Results are representative of three independent experiments. **(C)** Relative RRM2B mRNA expression was quantified by RT-qPCR in TNNT3-depleted cells following transfection with TNNT3 siRNA and treatment with Osimertinib. Transcript levels were normalized to GAPDH. **(D)** Representative images from colony formation assays in PC-9 and HCC827 cells transfected with control siRNA, RRM2B siRNA, or FLAG-tagged RRM2B, and treated with varying concentrations of Osi. **(E)** Quantification of colony density in treated cells from panel C. Data reflect the impact of RRM2B modulation on cellular resistance to Osi.

To investigate the role of RRM2B induction in the development of resistance to osimertinib, we performed colony formation assays in EGFR-mutant NSCLC cell lines PC-9 and HCC827. As shown in **Fig. 5D**, treatment with 2.5 nM osimertinib alone moderately reduced colony formation in control cells. However, co-treatment with osimertinib and RRM2B knockdown led to a marked reduction in both colony number and size, indicating increased sensitivity. Notably, the combination of 10 nM osimertinib and RRM2B depletion resulted in near-complete loss of colony-forming ability in both cell lines. In contrast, exogenous overexpression of RRM2B significantly enhanced colony formation, even in the presence of osimertinib, supporting its role in promoting long-term survival under EGFR inhibition. Quantification of colony intensity (**Fig. 5E**) confirmed that RRM2B knockdown significantly impairs clonogenic growth under osimertinib treatment, whereas cells with intact or overexpressed RRM2B retain higher proliferative capacity. These findings underscore the functional importance of RRM2B in maintaining replication fitness and genomic stability during EGFR-targeted therapy. While osimertinib suppresses canonical RRM2-dependent nucleotide synthesis, RRM2B upregulation emerges as a critical adaptive mechanism, facilitating resistance by restoring dNTP pools and supporting DNA replication under drug-induced stress.

### Activation of the CHK2 Pathway Following Osimertinib Treatment Is Important for Maintaining Resistance to Osimertinib

To investigate signaling pathways activated in response to osimertinib-induced DNA double-strand breaks (DSBs), we performed immunoblotting for key DNA damage response proteins in PC-9 and HCC827 cells. WB analysis revealed a pronounced suppression of CHK1 signaling, accompanied by concurrent activation of CHK2 (**Fig. 6A**). Osimertinib treatment led to a significant reduction in both CHK1 phosphorylation and total CHK1 protein levels, indicating robust downregulation of CHK1 activity. This decrease was not reversed by the proteasome inhibitor MG132 (**Fig. 6B**), suggesting that CHK1 suppression occurs via a proteasome-independent mechanism, likely involving reduced CHK1 transcription (**Supplementary Fig. S6**). In contrast, osimertinib treatment markedly increased CHK2 mRNA expression (**Supplementary Fig. S6**). This reciprocal regulation—loss of CHK1 activity alongside CHK2 activation—was consistently observed across cell lines and experimental replicates, highlighting a compensatory shift in DNA damage signaling under osimertinib-induced stress. These findings suggest that CHK2 is upregulated as a compensatory response to the suppression of CHK1 in the context of osimertinib-induced replication stress. To assess the functional relevance of CHK2 activation, we co-treated cells with osimertinib and the CHK2 inhibitor PV1019. Inhibition of CHK2 significantly sensitized both PC-9 and HCC827 cells to osimertinib (**Fig. 6C**), supporting a critical role for CHK2 in mitigating osimertinib-induced DNA replication stress. These results indicate that CHK2-mediated DNA damage signaling may serve as a protective, pro-survival mechanism under EGFR inhibition.

**Figure 6.**
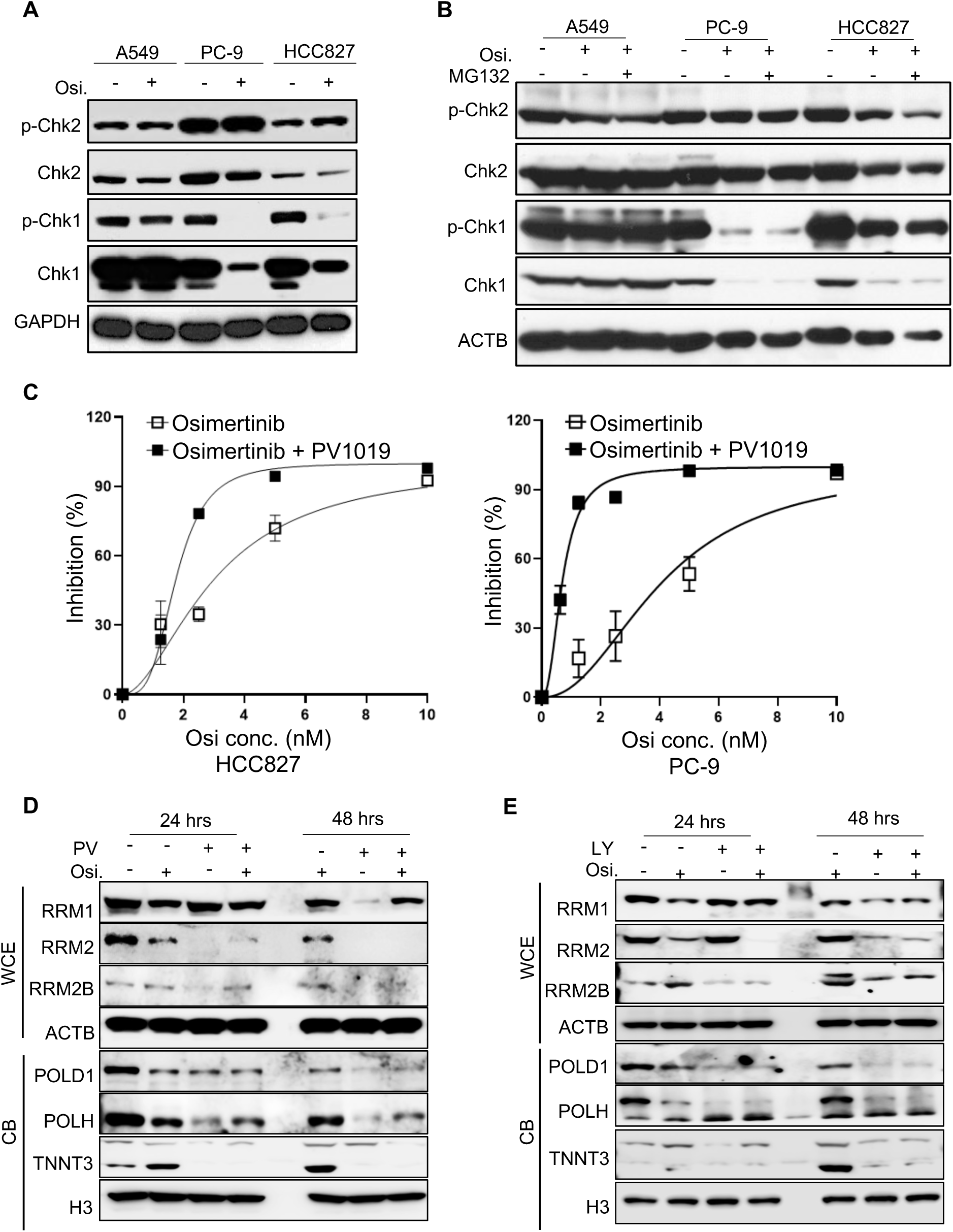
CHK2 pathway activation supports maintenance of osimertinib resistance. **(A)** A549, PC-9, and HCC827 cells were treated with 5 nM osimertinib (Osi). Protein levels of phosphorylated CHK1 (p-CHK1), phosphorylated CHK2 (p-CHK2), total CHK1, and CHK2 were analyzed by western blotting. GAPDH served as a loading control. Data are representative of three independent experiments. **(B)** Cells were treated as in panel A, with MG132 added 2 hours before harvest. Protein levels of p-CHK1, p-CHK2, CHK1, and CHK2 were assessed by western blotting. ACTB was used as a loading control. Data represent three independent experiments. **(C)** Proliferation assays in HCC827 and PC-9 cells treated with increasing concentrations of osimertinib alone or in combination with PV1019 (CHK2 inhibitor, concentration specified). **(D)** PC-9 cells were treated with DMSO, 5 nM Osi, 10 µM PV1019, or the combination for the indicated times. Nuclear fractions were analyzed by western blotting for RRM1, RRM2, RRM2B, POLD1, POLH, and TNNT3. ACTB and Histone H3 were used as loading controls. **(E)** PC-9 cells were treated with DMSO, 5 nM Osi, 10 nM LY2606368 (CHK1/CHK2 inhibitor), or the combination for the indicated times. Nuclear extracts were analyzed as in panel D. Data are representative of three independent experiments.

Further immunoblotting analyses revealed that inhibition of CHK2 impaired replication stress responses by reducing nuclear translocation of TNNT3 and transcriptional activation of RRM2B, thereby disrupting the compensatory restoration of RRM2 expression. As a result, both POLD1 and POLH levels in the chromatin fraction were significantly reduced in cells treated with either osimertinib alone or in combination with CHK2 inhibition (**Fig. 6D**). Similarly, combined inhibition of CHK1/2 using LY2606368 produced comparable effects, leading to depletion of both RRM2 and RRM2B and further impairing DNA replication (**Fig. 6E**). The reduction of TNNT3 in the chromatin fraction following treatment with osimertinib and CHK1/2 inhibitors may result from proteasome-mediated degradation of TNNT3 (**Supplementary Fig. 7A and B**), which subsequently leads to decreased transcription of RRM2B (**Supplementary Fig. 7C and D**). Collectively, these findings highlight a compensatory interplay between CHK1 and CHK2 pathways. Upon osimertinib-induced suppression of CHK1, CHK2 is activated to support DNA replication and repair through the regulation of RRM2B and polymerase. Disruption of this compensatory axis compromises replication integrity and sensitizes cells to EGFR-targeted therapy, underscoring CHK2’s potential role in promoting resistance to osimertinib.

### CHK1/2 Inhibition Delays the Development of Osimertinib Resistance and Enhances Therapeutic Efficacy in EGFR-Mutant NSCLC

To assess whether CHK1/2 inhibition affects the emergence of resistance to osimertinib, we conducted a stepwise drug escalation assay in EGFR-mutant NSCLC cell lines HCC827 and PC-9. Cells were continuously exposed to increasing concentrations of osimertinib—either alone or in combination with the CHK2 inhibitor PV1019 or the dual CHK1/2 inhibitor LY2606368—starting at 10 nM and escalating to 1000 nM over the indicated time course. As shown in **Fig. 7A and B**, both PC-9 and HCC827 cells acquired resistance to osimertinib monotherapy within approximately 120 to 200 days. In contrast, co-treatment with either PV1019 or LY2606368 significantly delayed the development of resistance in both cell lines. These results suggest that CHK1/2 pathway activation contributes to the adaptive resistance process and that inhibition of these checkpoint kinases impairs the cellular capacity to adapt to sustained EGFR inhibition.

**Figure 7.**
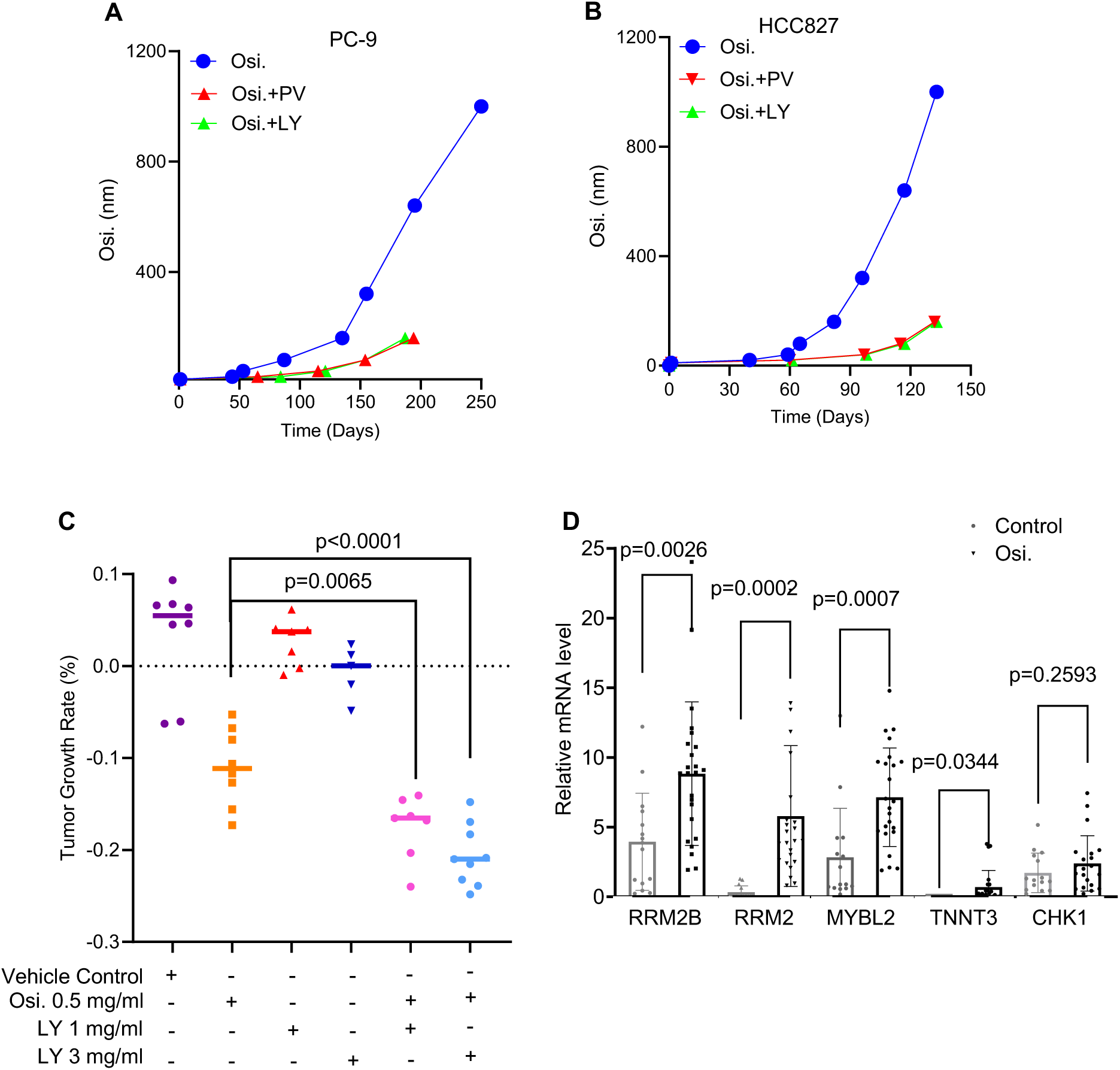
CHK1/2 inhibitor LY2606368 synergizes with osimertinib to suppress HCC827 tumor growth *in vivo*. Stepwise drug escalation assay to generate resistance to combined treatment with cell cycle checkpoint inhibitors (PV1019 or LY2606368) and osimertinib in **(A)** PC-9 and **(B)** HCC827 and cells. Cells were exposed to increasing concentrations of osimertinib, starting at 10 nM and escalating to 1000 nM over several weeks. At each step, proliferating cells were expanded until resistance was confirmed by sustained growth in 1000 nM osimertinib, assessed via trypan blue exclusion assay. **(C)** Tumor growth rates of HCC827 xenografts treated with DMSO (vehicle control, n=8), osimertinib (n=8), LY2606368 at 1 mg/kg or 3 mg/kg (n=5 each), or a combination of osimertinib with LY2606368 at each dose (n=7 and n=9, respectively). Tumor volumes were measured twice weekly and are presented as mean ± SEM. Statistical significance was determined using Student’s *t*-test. **(D)** Relative mRNA expression levels of RRM2B, RRM2, MYBL2, TNNT3, and CHK1 in tumor tissues following treatment with DMSO or osimertinib, as measured by quantitative RT-PCR.

To validate these findings *in vivo*, we established cell-derived xenograft (CDX) models using HCC827 cells in immunocompromised mice. Tumor-bearing mice were treated with vehicle, osimertinib alone, CHK1/2 inhibitor alone, or the combination. Tumor growth rate measurements revealed that the combination therapy induced the most pronounced tumor regression, significantly outperforming either agent alone (**Fig. 7C**).

To investigate molecular features associated with acquired osimertinib resistance, we performed gene expression profiling of resistant tumors. Analysis revealed significant upregulation of RRM2B, RRM2, and MYBL2 following osimertinib treatment (**Fig. 7D**), suggesting that restoration of dNTP pool balance is essential for the acquisition of resistance. Collectively, these results demonstrate that combined CHK1/2 inhibition delays cell adaptation to osimertinib-mediated RRM2 downregulation and enhances antitumor efficacy in both *in vitro* and *in vivo* models. These findings support a therapeutic strategy of co-targeting the DNA damage response to prevent or overcome resistance in EGFR-mutant NSCLC.

## DISCUSSION

This study identifies a critical role for dNTP metabolism in shaping the response of EGFR-mutant NSCLC cells to EGFR tyrosine kinase inhibition (TKI). We reveal a coordinated transcriptional and checkpoint signaling network that maintains dNTP pool balance and replication integrity under osimertinib treatment. Our findings delineate a dual regulatory axis—EGFR–MYBL2–RRM2 and CHK2–TNNT3–RRM2B—that enables cancer cells to adapt to TKI-induced replication stress. Disruption of this network impairs dNTP recovery, exacerbates DNA damage, and delays the emergence of osimertinib resistance, highlighting nucleotide metabolism as a tractable vulnerability in EGFR-driven NSCLC.

We demonstrate that osimertinib suppresses RRM2 expression by downregulating MYBL2, a transcription factor positively regulated by EGFR signaling. A major insight from this work is the impact of osimertinib treatment on pyrimidine dNTP homeostasis. Osimertinib significantly downregulates RRM2, a key regulator of dCTP and dTTP production, leading to the selective depletion of pyrimidine dNTP pools. This imbalance contributes to the slowing of replication forks, increased DNA damage, and the activation of compensatory DNA damage response (DDR) pathways. Our DNA fiber assays and comet analyses confirm that osimertinib-induced fork stress correlates with reduced availability of dNTPs, while supplementation with exogenous dNTPs rescues replication dynamics and limits DNA breakage. This is consistent with prior studies linking high RRM2 expression to poor prognosis and drug resistance in NSCLC [26, 27, 29, 30], and underscores RRM2 as a downstream effector of EGFR oncogenic signaling; and also consistent with prior studies showing that dTTP and dCTP depletion sensitizes cells to replication stress and induces genome instability [36, 38–42].

To compensate for RRM2 loss, NSCLC cells activate an alternative nucleotide synthesis program centered on RRM2B, a stress-inducible RNR subunit. We identify TNNT3 as a noncanonical transcriptional activator that translocates and accumulates in the nucleus upon CHK2 activation, driving RRM2B expression in p53-deficient contexts and enabling partial restoration of dNTP pools. Loss of TNNT3 or RRM2B exacerbates pyrimidine imbalance, increases DNA damage, and enhances osimertinib sensitivity, indicating that pyrimidine dNTP salvage via RRM2B is a key mechanism of drug resistance. TNNT3 has not previously been implicated in nucleotide metabolism or DNA replication, and our results highlight its functional relevance as a mediator of replication resilience.

Furthermore, we demonstrate that pharmacological inhibition of CHK2, or dual inhibition of CHK1/2, blocks RRM2B induction, thereby disrupting dNTP recovery and intensifying replication stress. Co-treatment with CHK1/2 inhibitors not only enhances the cytotoxicity of osimertinib but also significantly delays the acquisition of resistance both *in vitro* and in xenograft models. These findings highlight the therapeutic potential of combining EGFR inhibition with agents that target replication stress response pathways to exploit the metabolic fragility imposed by TKI-induced pyrimidine dNTP depletion; particularly in tumors with compromised p53 function that rely heavily on alternative stress-response pathways.

In conclusion, our study uncovers a previously unrecognized transcriptional and checkpoint network that preserves dNTP homeostasis and facilitates replication recovery under EGFR-targeted therapy. By identifying MYBL2, RRM2, TNNT3, and RRM2B as key effectors in this adaptive response and CHK2 as a potential therapeutic target, we propose a novel framework for understanding and disrupting drug resistance mechanisms in EGFR-mutant NSCLC. Targeting this adaptive rewiring through checkpoint kinase inhibition or nucleotide biosynthesis disruption may represent an effective strategy to enhance drug responses and delay resistance in EGFR-mutant NSCLC and improve outcomes for patients with advanced lung cancer.

## AUTHORS’ DISCLOSURES

B. Shen, L. Zheng, and T. Williams report grants from the NIH/NCI.

## AUTHORS’ CONTRIBUTIONS

Q. Xie: Data curation, formal analysis, investigation, visualization, manuscript writing and editing. Y.Y. Wang: Formal analysis, validation, investigation, visualization, manuscript writing and editing. Y. Lei: Formal analysis, validation, investigation, visualization, manuscript writing and editing. Z. Yao: Data curation, formal analysis, investigation, visualization, manuscript writing and editing. A. Fernandez: Formal analysis, validation, investigation, visualization, manuscript writing and editing. L. Yang: Investigation, formal analysis, supervision, funding acquisition, manuscript writing and editing. J.D. Hess: Formal analysis, visualization, manuscript writing and editing. T. Williams: Funding acquisition, formal analysis, manuscript review and editing., L. Zheng: Funding acquisition, formal analysis, investigation, methodology, manuscript review and editing., B. Shen: Conceptualization, formal analysis, supervision, funding acquisition, manuscript writing, review and editing. M. Li: Investigation, formal analysis, supervision, funding acquisition, manuscript writing and editing.

## ACKNOWLEDGMENTS

We thank Dr. Brian Armstrong at the COH Light Microscopy core facility for his technical support of AIRYSCAN confocal microscopy. This work was supported by NIH grants R01 CA085344 to B.H.S. and R50 CA211397 to L.Z. This work in the Li laboratory was supported by the University of South China’s startup funding to M.L. Research reported in this publication includes work performed using City of Hope shared resources supported by the National Cancer Institute of the National Institutes of Health under award number P30 CA033572.

## Notes

**CONFLICT OF INTEREST DISCLOSURES:** The authors declare no potential conflicts of interest.

### Competing Interest Statement

The authors have declared no competing interest.

