## Supplementary Figures and Tables for "Adaptive regulation of dNTP homeostasis confers osimertinib resistance in EGFR mutant non-small cell lung carcinoma"

### Supplementary Tables

**Table S1.** Oligonucleotides used in the current studies for RT-qPCR, ChIP-qPCR, siRNA, and quantitation of DNTP pools

### Supplementary Figures

**Figure S1.** Validation of siRNA knockdown efficiency.

**Figure S2.** MYBL2 regulates the sensitivity of PC-9 and HCC827 cells to osimertinib.

**Figure S3.** Complementing osimertinib-treated cells with dNTPs leads to the accumulation of DNA polymerase eta in the chromatin.

**Figure S4.** The induction of RRM2B expression following osimertinib treatment can be abolished by RRM2B siRNA transfection.

**Figure S5.** Induction of RRM2B expression following osimertinib treatment further induce DNA damage.

**Figure S6.** The induction of CHK2 expression following osimertinib treatment compensates for the reduction in CHK1 expression.

**Figure S7.** The activation of CHK2 pathway following osimertinib treatment stabilizes TNNT3 in chromatin, subsequently enhancing RRM2B expression.

**Supplementary Table S1. Oligonucleotides used in the current studies for RT-qPCR, ChIP-qPCR, siRNA, and quantitation of DNTP pools**

| RT-qPCR | |
|---|---|
| Target | Sequence (5'-3') |
| GAPDH-s | GTCTCCTCTGACTTCAACAGCG |
| GAPDH-as | ACCACCCTGTTGCTCTAGCCAA |
| RRM2-s | CTGGCTCAAGAAACGAGGACTG |
| RRM2-as | CTCTCCTCCGATGGTTTGTGTAC |
| RRM2B-s | ACTTCATCTCTCACATCTTAGCCT |
| RRM2B-as | AAACAGCGAGCCTCTGGAACCT |
| CHK1-s | CAACAAACCCCTCAAGAAAGG |
| CHK1-as | TGGATTGAATGTGCTTAGAAAATC |
| TNNT3-s | GAGGAGCAGTACGAAGAAGAAG |
| TNNT3-as | TGAGTTTGGGTCTCGGTTTC |
| MYBL2-s | CTTGAGCGAGTCCAAAGACTG |
| MYBL2-as | AGTTGGTCAGAAGACTTCCCT |
| ChIP-qPCR | |
| Target | Sequence (5'-3') |
| ACTINB promoter-FW | GCGTGACTGTTACCCTCAAA |
| ACTINB promoter-RV | GATGAAGGCTACAAACCTACCC |
| RRM2 promoter1-FW | GGCAAATCAGAAAGCCACATAG |
| RRM2 promoter1-RV | GTACTACTCATTGGGCGTCAA |
| RRM2 promoter2-FW | CTCAGCGGCCCTAACTTT |
| RRM2 promoter2-RV | CTTTCGATCCGTGTCCCT |
| siRNA | |
| Target | Sequence (5'-3') |
| EGFR siRNA-1# | GTGTGTAACGGAATAGGTATT (dTdT) |
| EGFR siRNA-2# | TCGTCAGCCTGAACATAACAT (dTdT) |
| EGFR siRNA-3# | GCTTAGGTGTTTGCTGAAAGT (dTdT) |
| MYBL2 siRNA-1# | CCACCACATCGAAGGAACA (dTdT) |
| MYBL2 siRNA-2# | GCCCAAGAGCACACCTGTTAA (dTdT) |
| RRM2B siRNA-1# | TGAGTTTGTAGCTGACAGATT (dTdT) |
| RRM2B siRNA-2# | GAGCTATTAGCTCCTCTAGAT (dTdT) |
| TNNT3 siRNA-1# | GCTCAACATCGATCACCTTGG (dTdT) |

| | |
|---|---|
| TNNT3 siRNA-2# | GGAAGTAGAGAGGCCAGAAAG (dTdT) |
| Quantitation of dNTPs | |
| dTTP detection template | TCGCTCGCTCTTGCCTCGGTCCTCGCTCGCTCTTGCCT CGGTCCTCGCTCGCTCTTGCCTCGGTCCTCGCTCGCTC TTGCCTCGGTCCTCGCTCGCTCTTGCCTCGGTCCTCGC TCGCTCTTGCCTCGGTCCTCGCTCGCTCTTGCCTCGGT CCTCGCTCGCTCTTGCCTCGGTCCTTTATTTGGCGGTG GAGGCGG |
| dATP detection template | AGACAGACACAAGACACAGACCAGACAGACACAAGA CACAGACCAGACAGACACAAGACACAGACCAGACAG ACACAAGACACAGACCAGACAGACACAAGACACAGA CCAGACAGACACAAGACACAGACCAGACAGACACAA GACACAGACCAGAGAGACACAACAGACGGAGGAAAT AAAGGCGGTGGAGGCGG |
| dCTP detection template | CCACTCACTCTTACCTCAATCCCCACTCACTCTTACCT CAATCCCCACTCACTCTTACCTCAATCCCCACTCACTC TTACCTCAATCCCCACTCACTCTTACCTCAATCCCCAC TCACTCTTACCTCAATCCCCACTCACTCTTACCTCAAT CCCCACTCACTCTTACCTCAATCCTTTGTTTGGCGGTG GAGGCGG |
| dGTP detection template | GGAGTGAGTGTGAGGTGAATGATGAGTGAGTGTGAGG TGAATGTAGAGTGAGTGTGAGGTGAATGATGAGTGAG TGTGAGGTGAATGTAGAGTGAGTGTGAGGTGAATGAT GAGTGAGTGTGAGGTGAATGTAGAGTGAGTGTGAGGT GAATGATGAGTGAGTGTGAGGTGAATGGTTTCTTTGG CGGTGGAGGCGG |
| Detection primer | CCGCCTCCACCGCC |

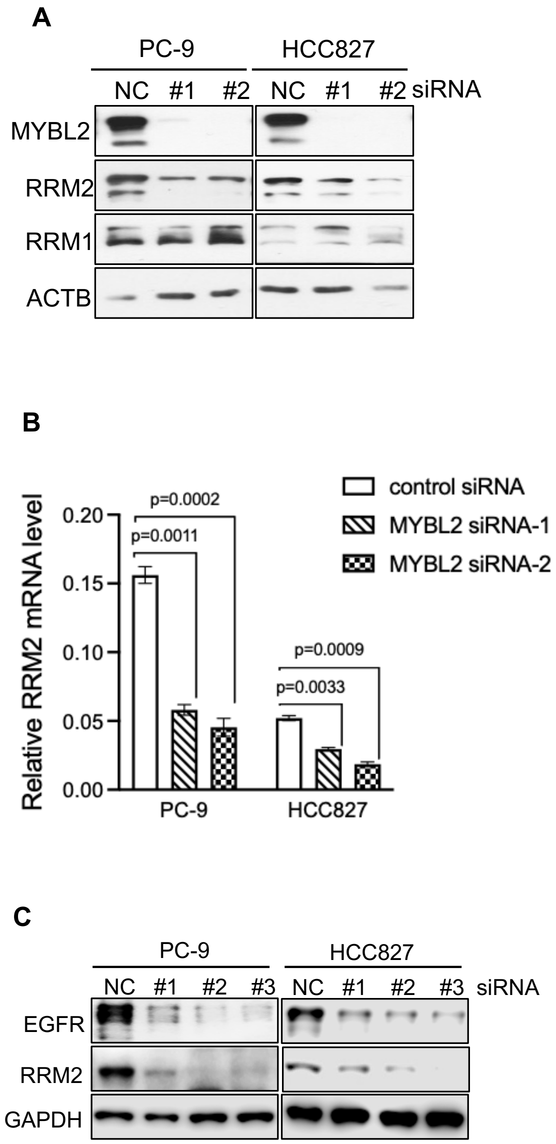


**Supplementary Figure S1**

**Supplementary Figure S1. Validation of siRNA knockdown efficiency.**

**(A)** PC-9 and HCC827 cells were transfected with control or MYBL2 siRNA. Protein levels of MYBL2, RRM2, and RRM1 were analyzed by western blotting. ACTB served as a loading control. Data are representative of three independent experiments. **(B)** RRM2 mRNA levels were measured by RT-qPCR in PC-9 and HCC827 cells following MYBL2 knockdown. Transcript levels were normalized to GAPDH. **(C)** PC-9 and HCC827 cells were transfected with control or EGFR siRNA. Protein levels of EGFR and RRM2 were examined by western blotting. GAPDH served as a loading control. Data are representative of three independent experiments.

**A**

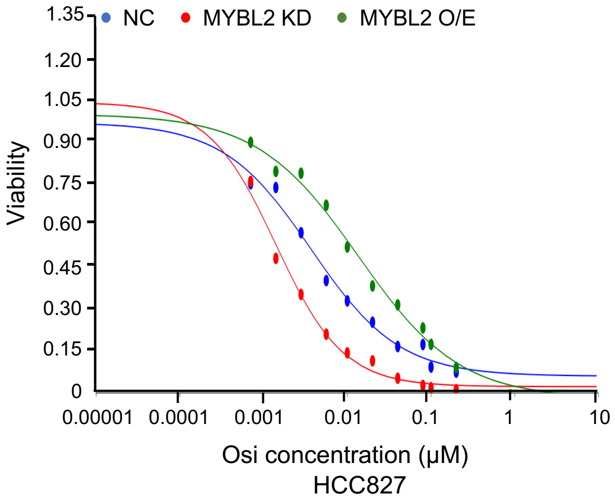


| Cell | $IC_{50}$ (nM) |
|---|---|
| HCC827 NC | 3.3 |
| HCC827 MYBL2 KD | 1.9 |
| HCC827 MYBL2 O/E | 10.2 |

**B**

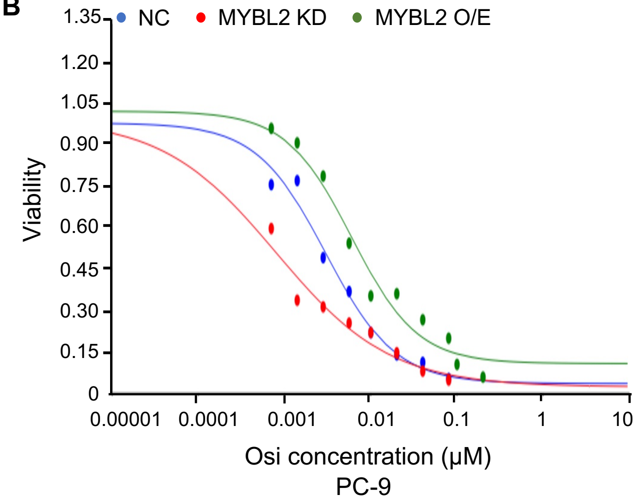


| Cell | $IC_{50}$ (nM) |
|---|---|
| PC-9 NC | 3.1 |
| PC-9 MYBL2 KD | 0.9 |
| PC-9 MYBL2 O/E | 6.0 |

**Supplementary Figure S2**

**Supplementary Figure S2. MYBL2 regulates the sensitivity of PC-9 and HCC827 cells to osimertinib.**

Cell proliferation inhibition assays were performed on **(A)** HCC827 and **(B)** PC-9 cells following MYBL2 knockdown or overexpression, in the presence of increasing concentrations of osimertinib.

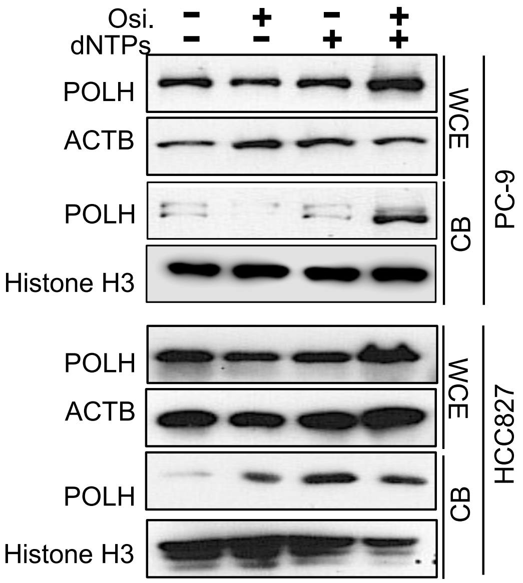


**Supplementary Figure S3**

**Supplementary Figure S3. Complementing osimertinib-treated cells with dNTPs leads to the accumulation of DNA polymerase eta in the chromatin.**

PC-9 and HCC827 cells were treated with DMSO, osimertinib (Osi), dNTPs, or their combination. Nuclear extracts were analyzed by western blotting for POLH. ACTB and Histone H3 were used as loading controls.

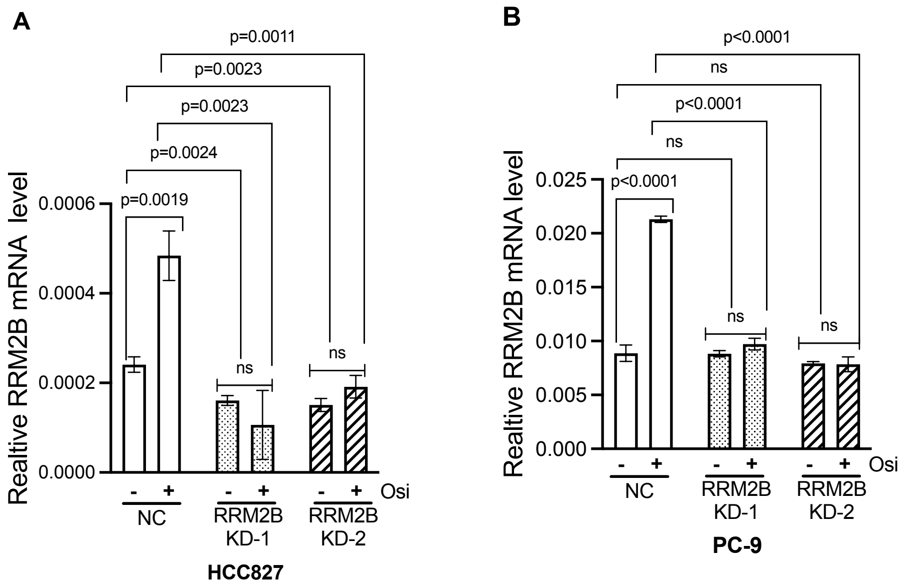


**Supplementary Figure S4**

**Supplementary Figure S4. The induction of RRM2B expression following osimertinib treatment can be abolished by RRM2B siRNA transfection.**

RRM2 mRNA expression was quantified by RT-qPCR in PC-9 **(A)** and HCC827 **(B)** cells following transfection with RRM2B siRNA and treatment with osimertinib. Transcript levels were normalized to GAPDH.

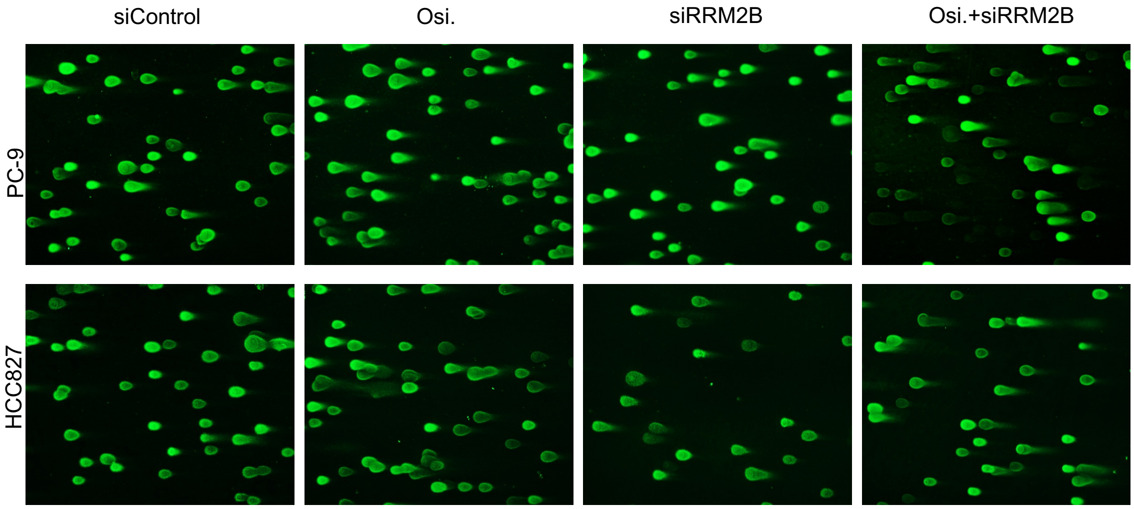


**Supplementary Figure S5**

**Supplementary Figure S5. Induction of RRM2B expression following osimertinib treatment further induce DNA damage.**

Representative comet assay images showing DNA damage in HCC827 and PC-9 cells following transfection with RRM2B siRNA and treatment with osimertinib (5 μM).

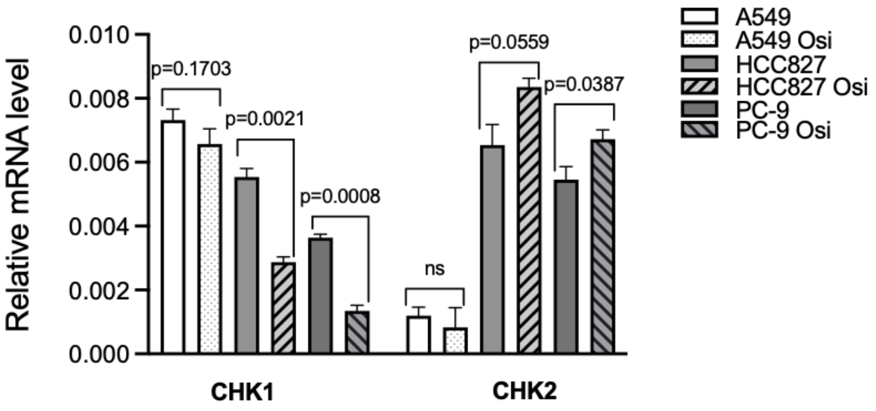


**Supplementary Figure S6**

**Supplementary Figure S6. The induction of CHK2 expression following osimertinib treatment compensates for the reduction in CHK1 expression.**

CHK1 and CHK2 mRNA levels were assessed by RT-qPCR in A549, PC-9, and HCC827 cells with or without osimertinib treatment. Transcript levels were normalized to GAPDH.

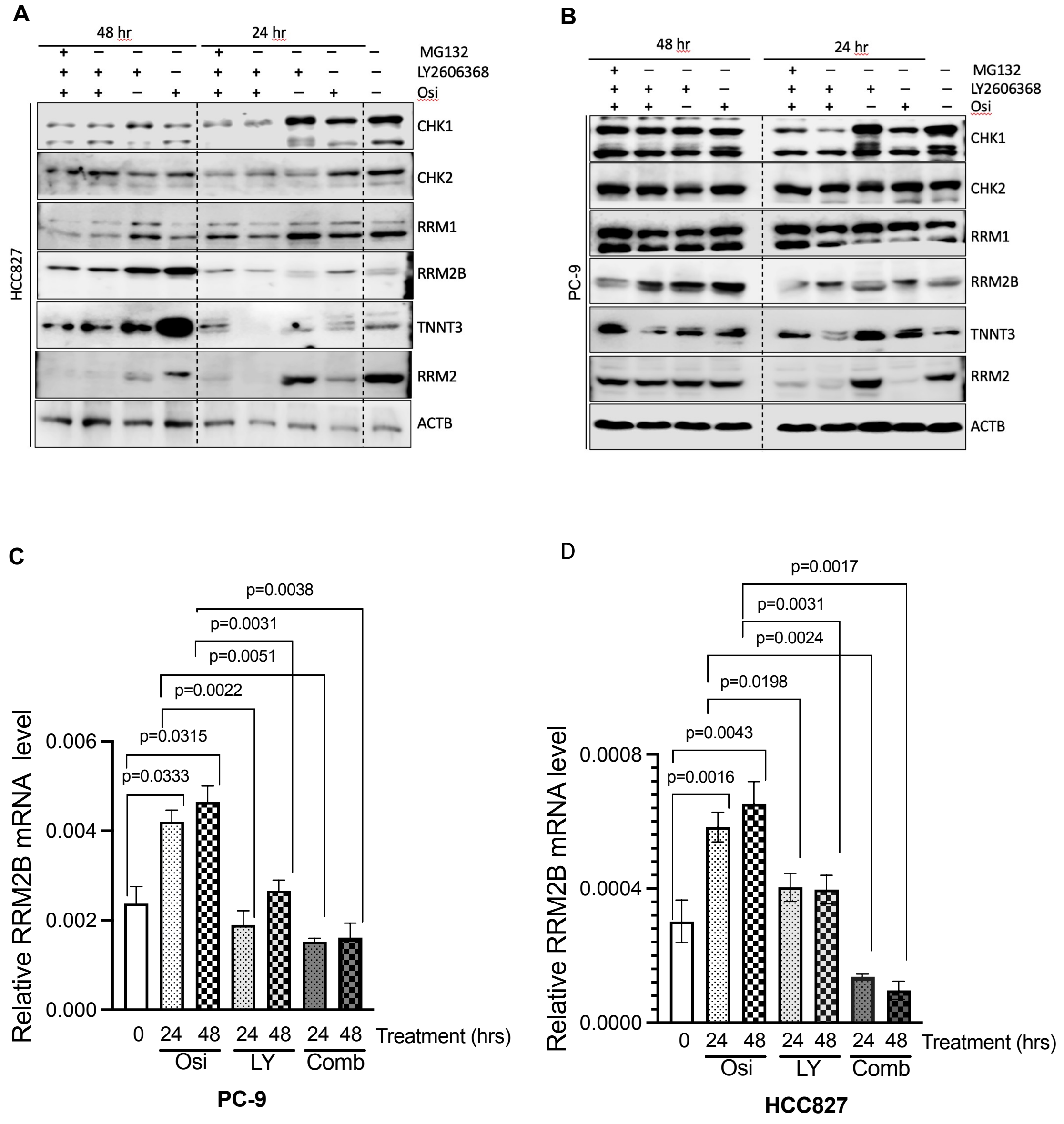


**Supplementary Figure S7**

**Supplementary Figure S7. The activation of CHK2 pathway following osimertinib treatment stabilizes TNNT3 in chromatin, subsequently enhancing RRM2B expression.**

**(A)** HCC827 cells were treated with DMSO, osimertinib, LY2606368, or their combination. MG132 was added 2 hours before harvest. Protein levels of CHK1, CHK2, RRM1, RRM2, RRM2B, and TNNT3 were analyzed by western blotting. ACTB and Histone H3 served as loading controls. **(B)** PC-9 cells were treated as in panel A and analyzed similarly. RRM2 mRNA expression was measured by RT-qPCR in **(C)** PC-9 and **(D)** HCC827 cells following the indicated treatments. Transcript levels were normalized to GADPH.
